# A critical role for heme synthesis and succinate in the regulation of pluripotent states transitions

**DOI:** 10.1101/2022.03.10.483773

**Authors:** Damien Detraux, Marino Caruso, Louise Feller, Maude Fransolet, Sébastien Meurant, Julie Mathieu, Thierry Arnould, Patricia Renard

**Affiliations:** Laboratory of Biochemistry and Cell Biology (URBC), NAmur Research Institute for LIfe Sciences (NARILIS), University of Namur (UNamur), Namur, Belgium; Institute for Stem Cell and Regenerative Medicine, University of Washington, Seattle, WA 98109, USA; Department of Comparative Medicine, University of Washington, Seattle, WA 98195, USA; Mass Spectrometry Facility (MaSUN), University of Namur, Namur, Belgium

**Keywords:** 2C-like cells, naive-to-primed, metabolism, succinate

## Abstract

Using embryonic stem cells (ESCs) in regenerative medicine or in disease modeling requires a complete understanding of these cells. Two main distinct developmental states of ESCs have been stabilized *in vitro*, a naïve pre-implantation stage and a primed post-implantation stage. Based on two recently published CRISPR-Cas9 knockout functional screens, we show here that the exit of the naïve state is impaired upon heme biosynthesis pathway blockade, linked to the incapacity to activate MAPK- and TGFβ-dependent signaling pathways. In addition, heme synthesis inhibition promotes the acquisition of 2 cell-like cells in a heme-independent manner caused by a mitochondrial succinate accumulation and leakage out of the cell. We further demonstrate that extra-cellular succinate acts as a paracrine/autocrine signal, able to trigger the 2C-like reprogramming through the activation of its plasma membrane receptor, SUCNR1. Overall, this study unveils a new mechanism underlying the maintenance of pluripotency under the control of heme synthesis.

## Introduction

The development and regeneration of an organism are two processes held by stem cells. These cells possess unique features such as an unlimited capacity for self-renewal and the ability to differentiate into various cell types. With their pluripotent phenotype, embryonic stem cells (ESCs) retain the ability to differentiate into all cell types of the embryo. First in mouse ^1,2^, then later in human ^3^, two main states of pluripotent stem cells have been described: the naïve ESCs, resembling the inner cell mass (ICM) of the pre-implantation embryo, and the primed ESCs, mirroring the epiblast of the post-implantation stage. While these cells represent timely close stages in embryo development, they display dramatic differences, such as developmental potential ^4,5^, epigenetic landscape, X-chromosome inactivation pattern and metabolic activity ^6–12^. Aside from these two pluripotent stages, ESC culture is known to be very heterogenous in terms of pluripotent or epigenetic marker expression ^13^. Interestingly, a small population (representing about 1 %) of mouse ESCs grown in naïve conditions displays features of the 2-cell stage embryo (2C-like population or 2CLCs) exhibiting extended potential ^14^. This subset also displays the expression of 2C-specific genes such as the *Zscan4* cluster, the retro-transposable element *MuERVL* or the master regulator DUX ^14–16^, a disappearance of chromocenters and loss of the core pluripotency protein OCT4 ^14^. However, despite the well-known differences between the different pluripotent states, little is understood about the molecular mechanisms governing the transition between them.

Several studies have started to address the question of the naïve-to-primed ESC transition using different screening methods to identify genes controlling this transition (reviewed in ^17^). Notably, a few studies have revealed mTORC1/2 as a critical component for the naïve-to-primed transition ^18–20^ that regulates key developmental pathways such as Wnt signaling ^6,18,21,22^. Among the hits of the different screens, we observed the recurrence of genes involved in the heme biosynthesis pathway. Although largely studied in hematopoietic stem cells, the roles of this biosynthetic pathway and this metabolite have never been studied in the context of pluripotency. In this study, we demonstrate the incapacity of mouse ESCs (mESCs) to exit from naïve toward the primed state under heme synthesis inhibition, a process caused by the inability to activate the core MAPK and TGFβ-SMADs signaling pathways. Unexpectedly, we also show that heme synthesis inhibition in naïve mESCs favors the emergence of 2CLCs in the cell population. Interestingly, this effect is heme-independent as hemin supplementation does not prevent it. We next demonstrate that the reprogramming in 2C-like state upon heme biosynthesis inhibition is actually caused by the accumulation of extra-mitochondrial succinate derived from the heme synthesis inhibition, acting as both paracrine and autocrine signals.

## Results

### Murine ESCs are dependent on heme biosynthesis to properly transition to the primed stage

In order to unveil the pathways required for the naïve-to-primed ESC transition, we compared the results of whole genome CRISPR-Cas9 screens previously published for mouse and human ESCs ^18,19^. The significant hits (*p*val< 0.05) were submitted to the DAVID functional annotation tools. For the human screen, positive hits for apoptosis were removed. Indeed, since a negative selection was applied to induce the death of primed human ESCs (hESCs), cells that acquired a resistance to cell death by mutating genes involved in apoptosis would be spared. **Figure 1a** displays the top Gene Ontologies (GO) for the DAVID biological processes showing “heme biosynthetic process” as one of the most enriched in both studies^18,19^. Seven out of 8 enzymes of this metabolic pathway (ALAD, PBGD, UROS, UROD, CPOX, PPOX and FECH) came out as positive hits in the CRISPR screen during the naïve-to-primed mESC transition. In hESCs, only the 4 cytosolic enzymes (PBGD, UROS, UROD and CPOX) were highlighted. Together, the results stress the importance of this metabolic pathway for the naïve-to-primed transition, although its role in non-hematopoietic stem cells is poorly understood.

**Figure 1:**
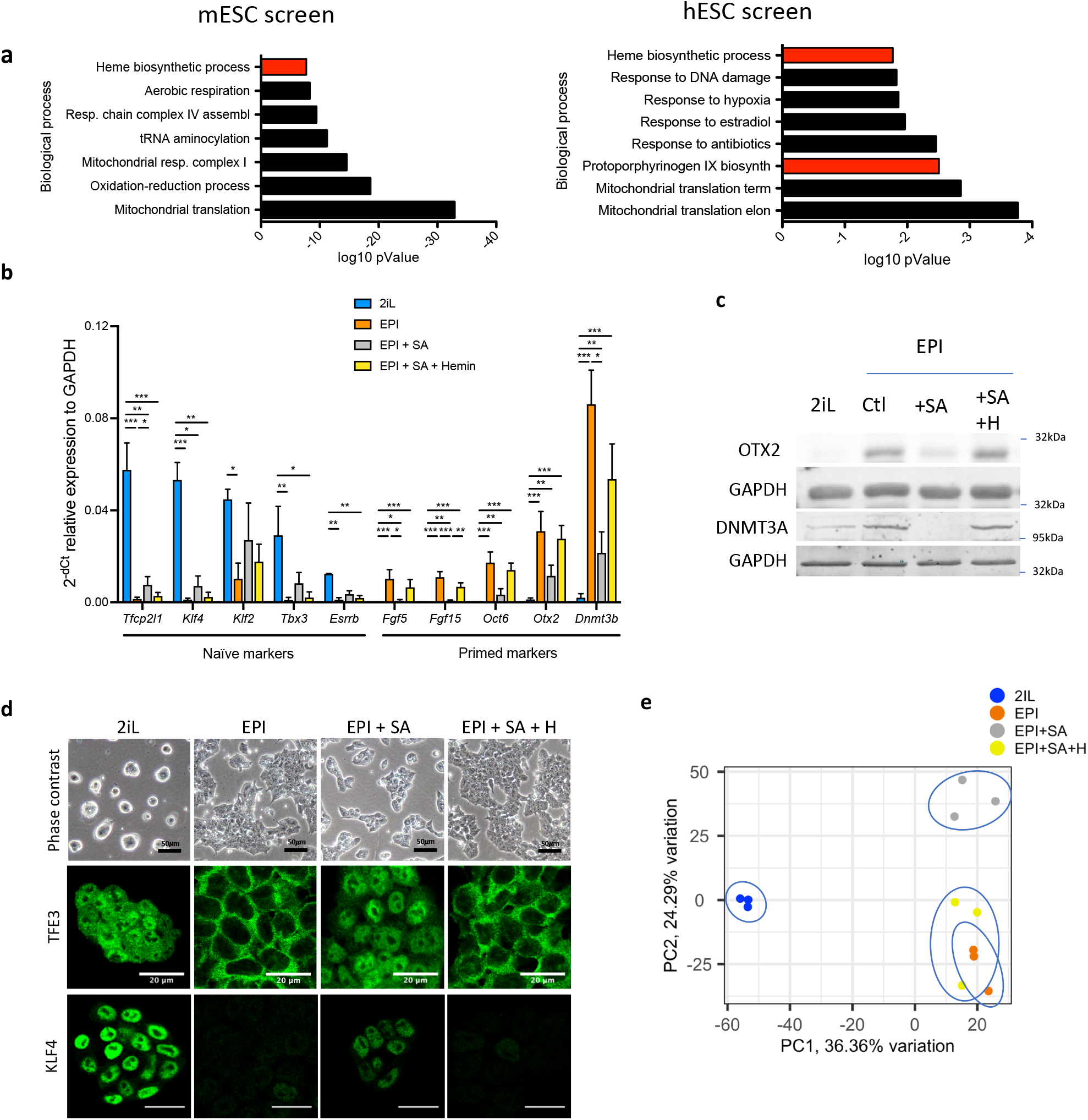
Heme synthesis inhibi4on impairs the exit of mESCs from the naïve state; effect mediated by heme. **a)** DAVID biological processes GO enrichment from two independent CRISPR-Cas9 screens for the naïve state exit, in mouse (leE panel)^17^ and in human (right panel)^18^. The heme biosyntheMc pathway is highlighted In red. **b)** Relative expression of naïve and primed markers of mESCs in naïve conditions (2iL), in transition for 2 days to the Epi stage (Epi) with or without 0.5 mM SA as heme synthesis inhibitor (Epi + SA 0.5mM) and 10 µM hemin supplementation (Epi + SA + Hemin 10 µM), assessed by RT-qPCR relative to GAPDH expression (*Tfcp2l1*, transcription factor CP2-like 1; *Esrrβ*, estrogen-related receptor β; *Klf2/4*, Kruppel-like factor 2/4; *Tbx3*, T-Box Transcription Factor 3; *Fgf5/15*, fibroblast growth factor-5/15; *Zic2*, zic family member 2; *Otx*2, homeobox protein 2). S.D. *p<0.05, **p < 0.01, ***p < 0.001. ANOVA-1. n=3 independent biological replicates. **c)** Western blot analysis of the protein abundance of OTX2 and DNMT3A relative to GAPDH as a loading control for cells in naïve conditions (2iL), in transition for 2 days to the Epi stage (EPI Ctl) with 0.5 mM SA (+SA) and 10 µM hemin (+H) supplementation. Representative blot of 3 biological replicates. **d)** Phase contrast micrographs of cells in naïve (2iL), or in transition to the primed (EPI) state with treatment with SA and hemin (H). Scale bar=50 μm. Confocal micrographs of mESCs in naïve stage or in transition for TFE3 (Transcription Factor Binding To IGHM Enhancer 3) and KLF4, in green. Scale bar =20 μm. TFE3 n=3 and KLF4 n=3 biological replicates **E)** Principal component analysis (PCA) of the normalized RNAseq data transcripts.

Experimentally, to trigger the exit of the naïve mESC state, the serum-free media with naïve cytokines cocktail (LIF; CHIR99021 and PD0325901) is switched to a media with fibroblast growth factor 2 (FGF2) and activin A for 48 h, allowing mESCs to gain post-implantation features. Using succinylacetone (SA) to inhibit heme synthesis, by interfering with ALAD activity ^23^, and 10 μM of hemin for its rescue, we show that, when mESCs are pushed for 48 h to exit the naïve stage (EPI) in the presence of SA (EPI + SA), the expression of the primed gene markers *Fgf5, Fgf15, Otx2, Oct6, Dnmt3a* and *Zic2* is prevented (**Fig. 1b**). In addition, the loss of expression for naïve markers (*Esrrb, Tfcp2l1, Klf2-4 and Tbx3*) is partially prevented. This was confirmed at the protein level by a decrease in the abundance of OTX2 and DNMT3A analyzed by western blot (**Fig. 1c**) and the increase in the abundance of KLF4 by immunofluorescence, when cells were treated with SA during the transition (**Fig. 1d)**. Furthermore, the subcellular localization of TFE3, mainly nuclear in naïve cells and only cytosolic in primed cells ^18,24^, remains nuclear in the presence of SA (**Fig. 1D**). Thus, hemin supplementation restores the gene expression, the protein abundance and the subcellular localization of TFE3 to levels similar to those found in cells incubated without SA (**Fig 1a-d**). Finally, principal component analysis (PCA) of the normalized gene expression from RNA sequencing also reveals the segregation of the cells treated with SA (EPI + SA) from either the controls (EPI) or the cells rescued with hemin (EPI + SA + H) (**Fig. 1e**). Overall, these results confirm the screens data by showing that the inhibition of heme biosynthesis impairs the naïve-to-primed mESC transition.

### Heme synthesis inhibition prevents the activation of key developmental signaling pathways

To identify the mechanisms involved in the failure to properly undergo the transition, we performed a gene set enrichment analysis (GSEA) with the KEGG pathways (Kyoto Encyclopedia of Genes and Genomes) between EPI and EPI + SA RNAseq data. Interestingly, many crucial signaling pathways involved in development are shown negatively enriched in SA-treated condition (**Fig. 2a**). We thus focused our attention on the pathways directly associated with the naïve-to-primed transition that is triggered by the combined presence of FGF2 and activin A in the growth culture media, especially since detailed analysis of the PC loadings driving the PC2 separation (**Fig. 1e**) highlights several SMAD pathway-related proteins (**Suppl. Fig. 1**). Strikingly, on the one hand, the MAPK-ERK1/2 pathway (downstream of FGF2) was not activated in EPI + SA cells, as shown by the absence of phosphorylation of ERK1/2 (**Fig. 2b**). On the other hand, the activin A-SMAD pathway activation was also compromised as shown by the difference in nuclear localization of SMAD2/3 when compared to the EPI cells (**Fig. 2c**). Interestingly, chemical inhibition of those two pathways by 1 μM PD0325901 (MEKi) and 5 μM SIS3 (TGFβ-SMADi) (**Fig. 2b-c**) shows that the inhibition of the SMAD translocation mimics the transition defects observed with SA as revealed by the gene expression analysis of naïve and primed markers (**Fig. 2d**). We thus conclude that a failure to activate these key cell signaling pathways, especially the activin A-SMAD pathway, impairs the transition to the primed stage, despite the presence of their respective ligands in the cell culture media.

**Figure 2:**
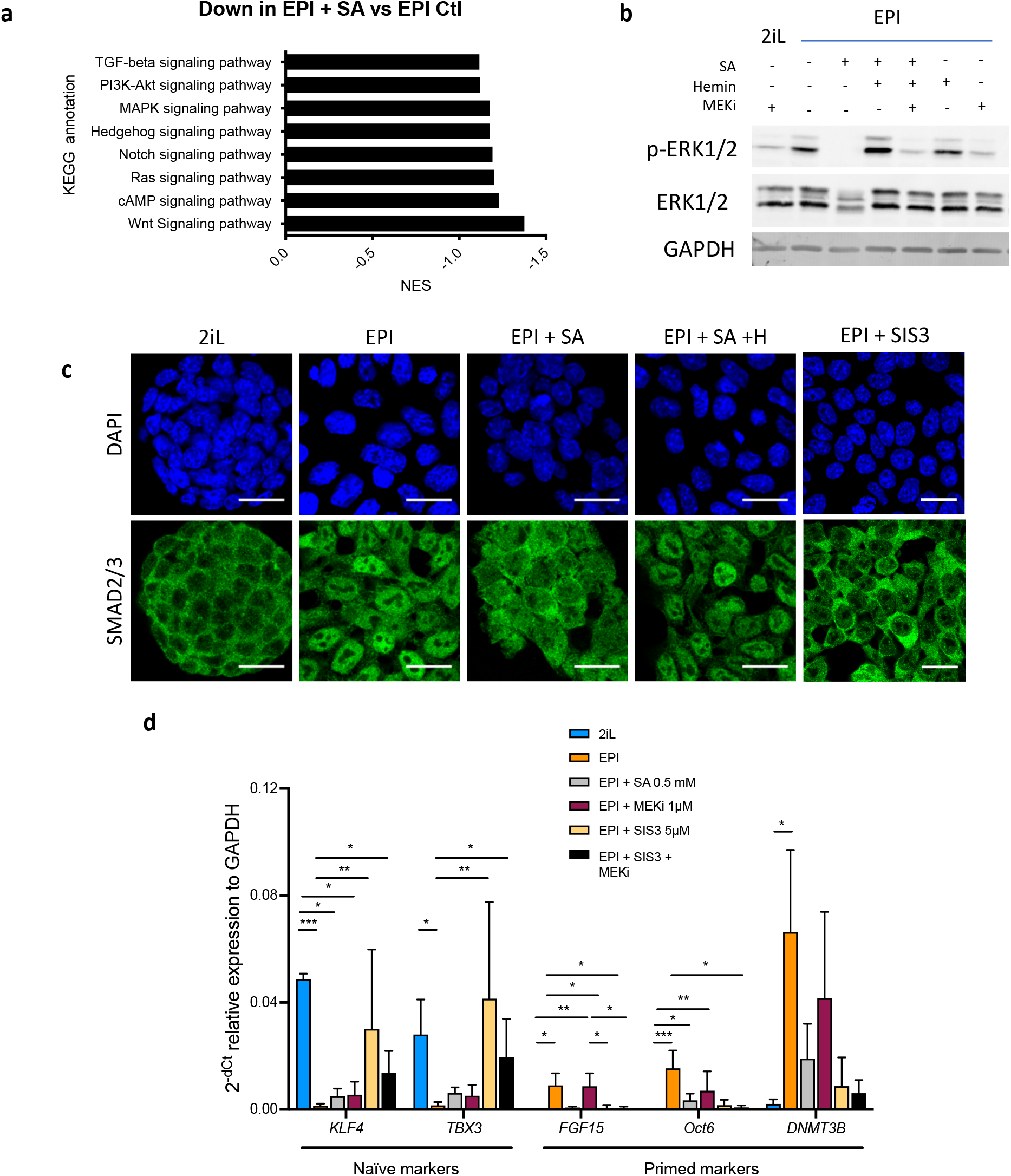
SA prevents the activation of the MAPK and Activin A-SMAD pathways during the mESC transition. **a)** GSEA performed on RNAseq data were analyzed for KEGG pathways. Down regulated KEGG pathways in EPI+SA versus EPI Ctl are represented as normalized enrichment scores (NES). **b)** Western blot analysis of the protein abundance of ERK1/2 and phospho-ERK1/2 (Thr202/Tyr204) relative to GAPDH as a loading control for cells in naïve conditions (2iL), in transition for 2 days to the Epi stage (EPI) with 0.5 mM SA, 10 µM hemin (H) supplementation and/or 1 µM MEK inhibitor (PD0325901; MEKi). Representative blot of 3 biological replicates. **c)** Confocal analysis of the immunostaining of SMAD2/3 (green) in cells in naïve conditions (2iL), in transition to the Epi stage (EPI) with SA, hemin (H) supplementation or a SMAD3 inhibitor (SIS3, 5 µM), representative of 3 independent experiments. **d)** Relative expression of naïve and primed markers of mESCs in naïve (2iL), in transition (Epi) with or without SA 0.5 mM, Hemin 10 µM, SIS3 5 µM, MEKi 1 µM (PD0325901), assessed by RT-qPCR relative to GAPDH expression. Data shown as mean +/-S.D. n=3 independent biological replicates.

### Blockade of heme synthesis in naïve mESCs triggers the activation of a 2C-like program

In addition to this defect in the exit from the naïve state, we found that treatment of naïve 2iL cells with SA also modifies the global gene expression as 2iL + SA samples cluster away from 2iL control cells when analyzed by PCA performed on RNAseq data (**Fig. 3a**). Our attention was thus drawn on markers reported to be expressed in the 2-cell embryo. Indeed, a small proportion of the mESC population actually expresses a gene signature reminiscent of the 2C stage, these cells thus called 2C-like cells (2CLCs)^14^. Indeed, the expression of the 142 genes previously identified as upregulated in this 2C-like population ^14^ was statistically upregulated in 2iL versus 2iL + SA (**Fig. 3b**) and this was further confirmed by RT-qPCR for a selection of the most common markers (**Fig. 3c**) and by western blot analysis of ZSCAN4 abundance (**Fig. 3c**). As the studies that characterize this 2C-like population report an important heterogeneity at the population level ^14,25^, we quantified the fraction of ZSCAN4^+^ and MUERVL-GAG^+^ cell populations by immunostaining followed by confocal microscopy observations. As shown in **Fig. 3e and f**, treatment with SA increases the fraction of 2C-like cells by 2 to 3 folds.

**Figure 3:**
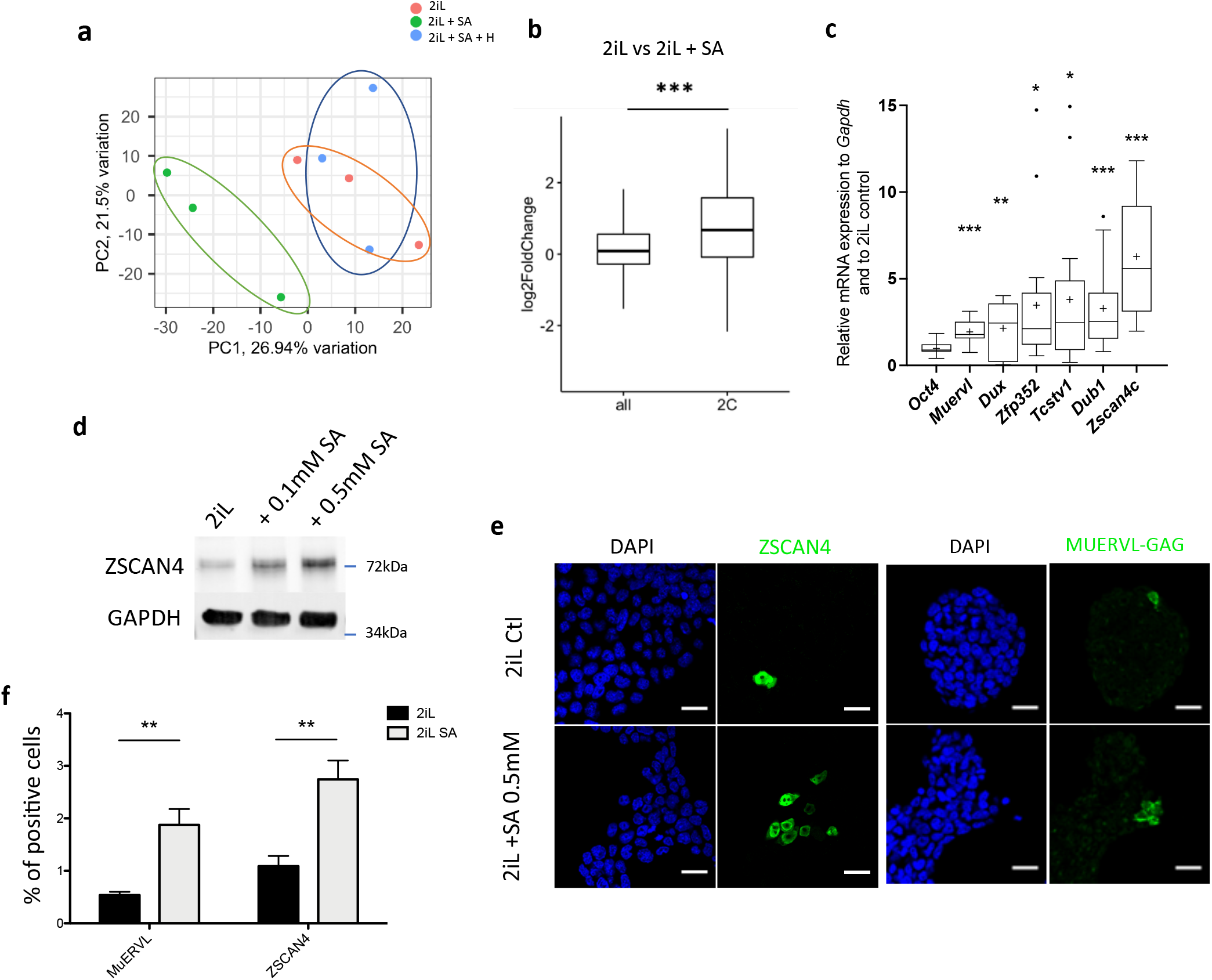
Heme synthesis inhibi1on pushes mESCs toward a 2C-like stage. **a)** Principal component analysis (PCA) of the normalized RNAseq data transcripts of naïve mESCs (2iL) treated for 48h with 0.5 mM SA ± 10 μM Hemin. **b)** Boxplot of mean Log2FC of 2C markers defined in ^14^ or all analyzed mRNAs in 2iL+SA versus 2iL control cells. Statistical significance is calculated by Student T-test. p<0.001. **d)** Relative expression of 2C gene markers of mESCs assessed by RT-qPCR relative to *Gapdh* expression and to 2iL naïve control represented as a Tukey box and whisker plot. The line represents the median and the + is the mean. *Oct4*= octamer-binding transcription factor 4, *Muervl*= murine endogenous retrovirus-like, *Dux*= double homeobox, *Zfp352*= Zinc-finger protein 352, *Tcstv1*= 2-cell-stage variable group member 1, *Dub1*= Ubiquitin Specific Peptidase 36, *Zscan4c*= Zinc Finger And SCAN Domain Containing 4, isoform c. n=9 **e)** Western blot analysis of ZSCAN4 protein abundance relative to GAPDH as a loading control in naïve mESCs (2iL) treated for 2 days with SA at 0.1 mM or 0.5 mM. Representative image of 3 independent biological replicates. **f)** Immunostaining of ZSCAN4 (green; led pannels) or MUERVL-GAG (green; right pannels) in untreated naïve (2iL control) or treated with SA at 0.5mM. DAPI is used as a nuclear counterstain. Scale bar= 20μm. **g)** Percentage of MUERVL-or ZSCAN4-positive cells in the whole population of naïve (2iL) mESCs or naïve treated or not with SA (2iL SA), counted from confocal micrographs as in (f) with 10 images per conditions for at least 1000 cells per condition. n=4 independent biological replicates. Results expressed as mean +/-S.D. ** p < 0.01 ; T-Tests.

As the 2C-like population is reminiscent of the totipotent state of stem cells, we next tested whether treatment with SA could increase the ability of mESCs to differentiate into trophoblast stem cells. To test this hypothesis, mESCs were first incubated for two days in the 2iL medium with or without SA. The medium was then switched to the trophoblast differentiation medium based on a combination of FGF4 and TGF-β1, as previously reported ^26^ (**Suppl. Fig. 2a**). After 6 days of trophoblast induction, three features were analyzed: cell morphology, gene expression and protein abundance of trophoblast markers. An increase in the abundance of cells with a different morphology was observed, showing enlargement of the cell surface, reminiscent of the giant cells derived from the trophoblast (**Suppl. Fig. 2b**) ^27,28^. This difference in phenotype is in line with the increase in expression of the trophoblast lineage markers *Eomes, Hand1, Id1* or *Elf5* (even if the differences are not statistically significant in our experimental conditions) and in the abundance of GATA3 (**Suppl. Fig. 2c-d**). Altogether, these findings suggest that a 48 h-treatment with SA is able to expand the potential of mESCs toward the extraembryonic lineage.

### The induction of the 2C-like program is succinate-dependent

Interestingly, and as opposed to the naïve-to-primed setup, the observed phenotype seems independent of heme as it is not rescued by hemin supplementation (**Fig. 4a-b**). Previous reports have shown that heme biosynthesis consumes a lot of the glycine and succinyl-CoA precursors in the mitochondria, acting as some sort of “succinyl-CoA sink” ^29^. We thus hypothesized that heme synthesis inhibition would increase the abundance of succinyl-CoA in mitochondria, that could then exit the organelle in the form of succinate and accumulate in other cell compartments. Furthermore, another report showed that the knock-down of succinate dehydrogenase subunit B (SDHB) expression could increase global protein succinylation throughout the cell, even affecting histone succinylation status, demonstrating the ability of succinate to exit mitochondria ^30^. We thus assessed the abundance of succinylated proteins in naïve cells treated or not with SA using a pan-succinyllysine antibody (**Fig. 4c**). In basal conditions, the bulk of succinyl-lysine modifications is located in the mitochondria (**Suppl. Fig. 3**), as expected. However, we observe a dramatic increase in the signal associated with succinylated proteins in all subcellular compartments when heme synthesis is inhibited, an effect that is not (or very partially) rescued upon hemin supplementation (**Fig. 4c**). Since this increase in protein succinylation in non-mitochondrial compartments involves the exit of succinate through the mitochondrial membranes, we postulated that blocking the exit of this metabolite from mitochondria would prevent the acquisition of widespread succinyl-lysine post-translational modifications and impair the acquisition of the 2C-like cells markers, only if this phenotype is dependent on increased succinate concentration. The inhibition of mitochondrial succinate exit was achieved using diethyl butylmalonate (BM), an inhibitor of SLC25A10, the succinate transporter in the inner mitochondrial membrane ^31^. As hypothesized, addition of BM combined to SA prevents most of the accumulation of succinyl-lysine signal (**Fig 4c**). This decrease in global lysine succinylation is correlated to a rescue of both the increase in 2C markers and the proportion of ZSCAN4 or MUERVL-positive cells in the population (**Fig. 4a and d**). Altogether, this shows that an accumulation of extra-mitochondrial succinate in naïve mESCs induces a 2C-like phenotype. In order to observe the levels of protein succinylation, and by extension the levels of succinate, in endogenous 2CLC of the mESC population, we took advantage of a previously described reporter cell line for this 2C-state, characterized by the stable insertion of a construct containing a turboGFP-coding gene under the control of the MUERVL long terminal repeat (2C:::turboGFP)^16,32^. The simultaneous observation of the endogenous GFP fluorescence, the absence of OCT4 ^14^ and the immunostaining of the succinyl-lysine residues showed an increase in protein succinylation specifically in the 2C-like cells (GFP+; OCT4-) sub-population (**Fig 4e-f**).

**Figure 4:**
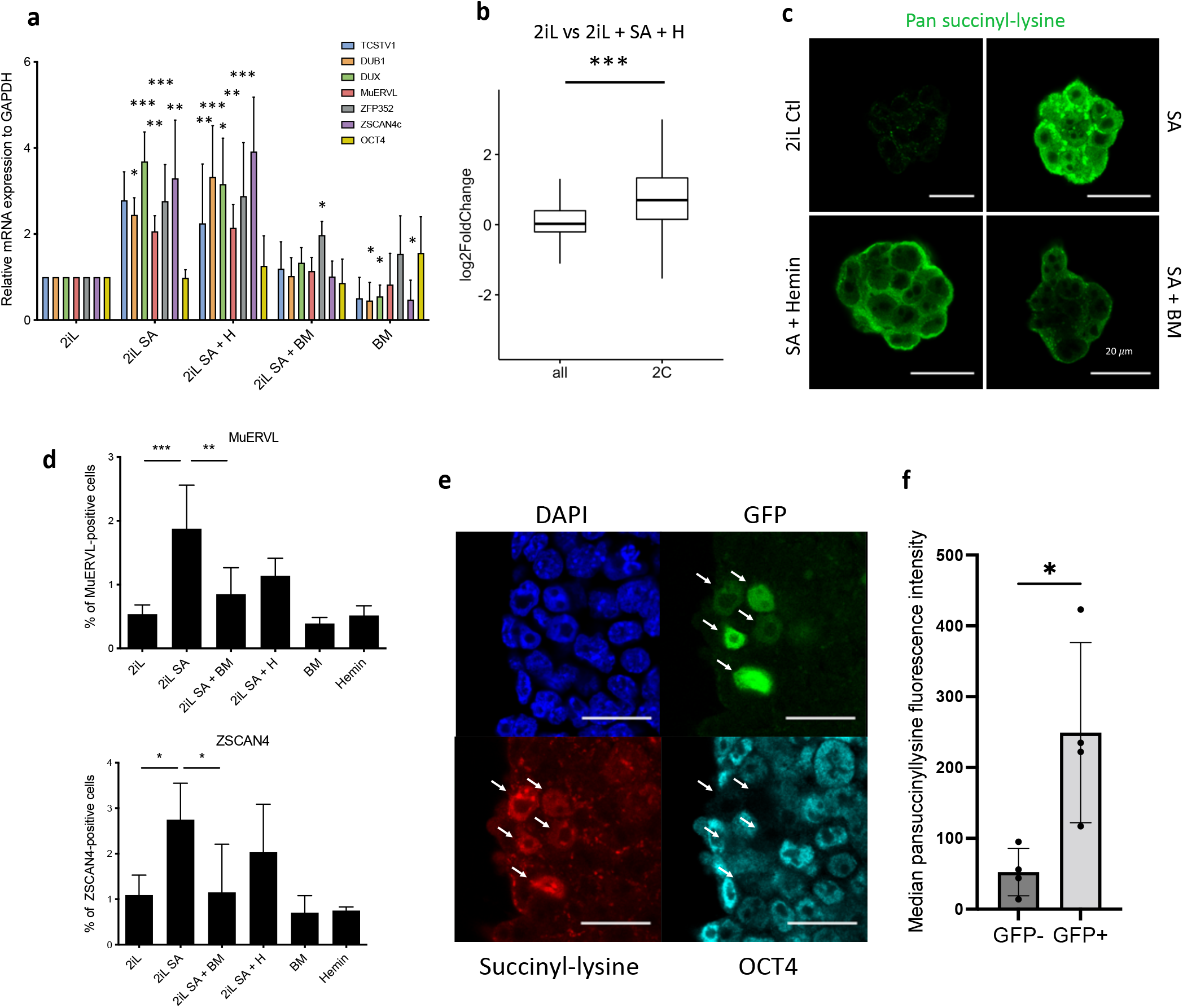
mESC “2C-like” reprogramming by SA is due to extramitochondrial succinate accumulation. **a)** Relative expression of 2C markers of mESCs assessed by RT-qPCR relative to GAPDH expression and normalized to 2iL naïve control, in mESCs treated with 0.5 mM SA (2iL SA), with or without 10 μM Hemin (2iL SA + H), 1 μM diethyl butylmalonate (2iL SA + BM) or BM 1μM alone (BM). S.D. * p < 0.05, **p < 0.01, ***p < 0.001. ANOVA-1. n=4 independent biological replicates. **b)** Boxplot of mean Log2FC of 2C markers defined in ^14^ or all analyzed mRNAs in 2iL+SA+H versus 2iL control cells. Statistical significance is calculated by Student T-test. p<0.001. **C)** Immunostaining of succinylated lysines (green) in mESCs treated with 0.5mM SA, with or without 10 μM Hemin (SA + Hemin) or 1uM diethyl butylmalonate (SA + BM 1μM). Representative image of n=3 independent experiments. Scale bar= 20μm. **d)** Percentage of MUERVL or ZSCAN4-positive cells in the whole population of naïve (2iL) mESCs or naïve treated with SA (2iL SA) with or without 10 μM hemin (2iL SA + H) or 1 μM diethyl butylmalonate (2iL SA + BM). n=4 independent biological replicates. Results expressed as mean +/-S.D. * p < 0.05, **p < 0.01, ***p < 0.001 ; ANOVA-1. **e)** Immunostaining of succinyl-lysine residues (Red) and Oct4 (cyan) of TBG4 cells (ES-E14TG2a mESCs with a 2C-GFP (green) reporter construct)^32^. Representative image of n=3 independent experiments. Scale bar= 20 μm. Arrows indicate 2CLCs according to the GFP reporter. **f)** Quantification of the median fluorescence intensity of the succinyl-lysine in the GFP+ and GFP-populations of TBG4 mESCs separated by flow cytometry. n=4 independent biological replicates. * p< 0.05

Since we showed that the increase of the reprogramming of some mESCs to a 2C-like state after heme synthesis inhibition was the result of succinate exit from mitochondria, we then aimed to further confirm these results, by inducing a mitochondrial accumulation of succinate, independently of heme biosynthesis inhibition. The SDH enzyme inhibition, known to provoke succinate accumulation ^30^, was achieved by exposing naïve 2iL mESCs to Atpenin A5 (AA5) ^33^. AA5 strikingly induces the protein lysine succinylation (**Fig. 5a**), an increase in the number of ZSCAN4-positive cells (**Fig. 5b**) and the expression of 2C markers (**Fig. 5c**). These effects were counteracted by the addition of BM to prevent the succinate exit from the mitochondria (**Fig. 5b**), confirming the involvement of succinate in the process.

**Figure 5:**
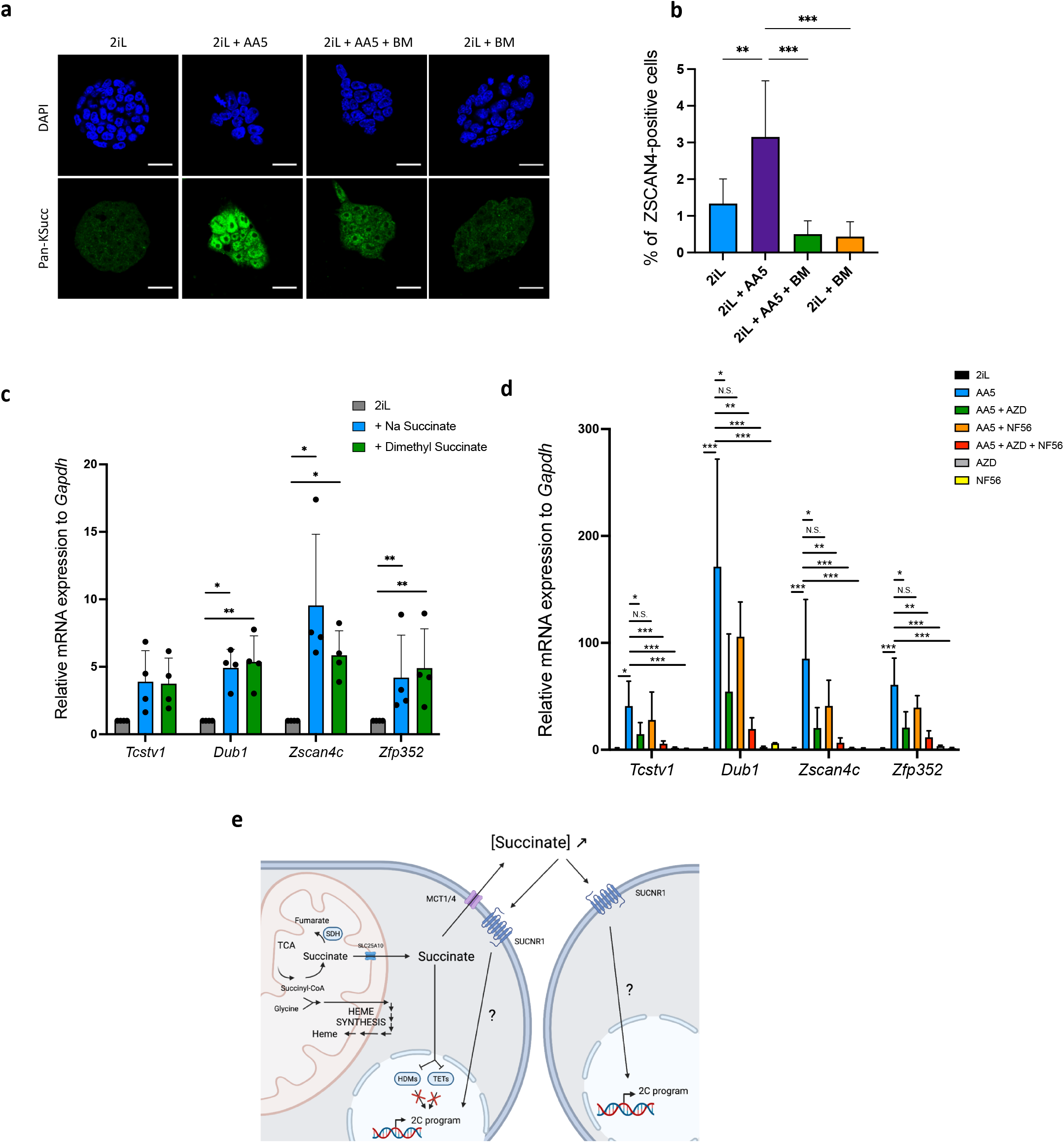
Inhibi-on of SDH leading to an increase in succinate accumula-on recapitulates the increase in 2CLCs due to a leakage to the extracellular space. **a)** Immunostaining of succinylated lysine residues (green) in mESCs treated with 250nM AA5, with or without 1uM diethyl butylmalonate (BM). Representative image of n=3 independent experiments. Scale bar= 20μm **b)** Percentage of ZSCAN4-posiEve cells in the whole population of naïve (2iL) mESCs or naïve treated with 250nM AA5, with or without 1μM diethyl butylmalonate (BM), counted from confocal micrographs with 10 images per conditions for at least 1000 cells per condition. n=3 independent biological replicates. **c-d)** Relative expression of 2C gene markers of mESCs assessed by RT-qPCR relative to *Gapdh* expression in naïve 2iL mESCs and treated with 16mM dimethyl succinate, 40mM sodium succinate (Na succinate), 1μM of MCT1 and 4 inhibitor (AZD3965; AZD) and 1μM SUCNR1 receptor antagonist (NF-56-EJ40; NF56). Data shown as mean +/-S.D. *p<0.05, **p < 0.01, ***p < 0.001. ANOVA-1 followed by a Tukey post-testusing AA5 as reference. n=4. **e)** Graphical representation of the succinate-induced 2CLC reprogramming. Created with Biorender.com.

### Succinate acts through the activation of its receptor, SUCNR1, to trigger the 2C-like program

Once in the cytosol, succinate could either inhibit the α-ketoglutarate (αKG)-dependent-dioxygenases or exit the cell through the monocarboxylate transporters (MCTs) and act as an extracellular signal molecule. Indeed, succinate is a reaction product of hydroxylation reactions catalyzed by αKG-dependent dioxygenases. Three families of dioxygenases are known to be sensitive to an increase in succinate concentration: the prolyl-hydroxylases (PDHs), the Ten-eleven translocation (TET) methylcytosine dioxygenases and the histone demethylases (HDM). Of these three classes, we ruled out a role for PHDs responsible for HIF1α stabilization as the stabilization of this transcription factor was neither induced by SA nor by AA5 treatment (**Suppl. Fig. 4**). Since recent reports in the literature have highlighted a crucial role of the epigenetic landscape and a role of the TET proteins in the acquisition of the 2CLC phenotype ^25,34–37^, we then focused our attention on the histone 3 (H3) and DNA methylation landscapes and observed a robust increase in the amount of trimethylation of lysines 4, 9 and 27 of H3 (H3K4,-K9, -K27) along with an increase in the global abundance of 5mC when mESCs were treated with either SA or AA5 (**Supp. Fig. 5d**). However, chemical inhibition of HDMs by JIB04 ^38^ and/or TET by TETin-C35 ^39^ does not trigger a rise in the proportion of 2CLCs or in 2C marker expression (**Suppl. Fig. 5e-f**). We then suspected that succinate could act instead as a paracrine signal after its release in the extracellular media. Indeed, exogenous supplementation with a membrane impermeable (sodium succinate) or permeable (dimethylsuccinate) form of succinate were able to increase the 2CLC marker expression (**Fig. 5c)**. We further found that inhibition of the succinate exit through the plasma membrane, by targeting MCT1 and 4 with the AZD3965 inhibitor ^40,41^ combined to inhibition of the succinate receptor SUCNR1 by NF-56-EJ40 ^42^ is able to bring down the rise in 2CLC marker expression triggered by AA5 (**Fig. 5d**). These findings strongly support succinate acting as a paracrine/autocrine signal. Together, these sets of data highlight the critical role of succinate in the acquisition and maintenance of pluri/totipotent states in mESCs by a totally new mechanism.

## Discussion

The comparison of two different genome-scale CRISPR screens in the exit of the naïve state of mouse and human ESCs ^18,19^ revealed the importance of heme biosynthesis for the transition to another pluripotent state, the primed state. Using succinyl acetone (SA), a specific inhibitor of ALAD that catalyzes the second reaction of the pathway, and hemin to rescue the heme defect, we first confirmed the requirement of this pathway for the naïve-to-primed transition in mESCs. This is in accordance with the embryonic lethality at the implantation stage of mouse embryos knockout for the heme synthesis pathway enzymes since the naïve-to-primed transition recapitulates, *in vitro*, this critical step (shown for HMBS ^43^, UROS ^44^, UROD ^45^, CPOX ^46^ and FECH ^47^). RNAseq analysis of naïve cells undergoing transition to the primed stage in the presence of SA with or without addition of hemin revealed a failure to properly activate the MAPK and TGFβ-SMAD pathways in response to heme biosynthesis inhibition. It has been previously demonstrated that the proper activation of these two pathways is required to proceed with the transition to the primed stage ^48–51^. Intriguingly, while the inhibition of SMAD3 was able to mimic the effect observed by heme synthesis inhibition, the inhibition of MAPK by MEK inhibition proved to be inefficient at blocking the process, in contradiction to its role in the progression of pluripotency ^48,49,51^. Such a connection between heme synthesis inhibition and signaling pathways has only been reported once in PC12 neuronal cells, with SA blocking the activation of the MAPK despite the presence of neuronal/nerve growth factor (NGF)^52^. Thus, it seems to be a conserved mechanism among various cell identities/types, as it is also observed in our study for cells responding to FGF2 signals. While the link between heme and these crucial signaling pathways remains unknown, it brings to light a crucial importance of this metabolite and its synthesis pathway, so far poorly understood, in the context of pluripotency.

Aside from its effect on the exit of the naïve stage, we showed here that the inhibition of heme synthesis also triggers a reprogramming toward a 2C-like stage. Indeed, mESCs cultured in 2iL conditions have been previously defined as a heterogenous population that naturally includes a small percentage of cells displaying features of the 2-cell stage embryo ^14,32^. Interestingly, this 2CLC population is transient and cycles back and forth to a naïve pluripotency state ^14,53^. Unexpectedly, heme synthesis inhibition favors this reprogramming as shown by the increase in 2C-like markers in the whole population and the proportion of ZSCAN4 or MUERVL-positive cells. Strikingly, this effect is clearly dependent on the accumulation of succinate as it is i) not rescued by hemin, ii) blocked by inhibition of the mitochondrial succinate transporter and iii) phenocopied by SDH inhibition. Additionally, MUERVL-positive cells spontaneously emerging in naïve ESC colonies endogenously display a high level of succinylated proteins, supporting a role for this metabolite in the identity of the 2C-like state. The role of several metabolites in the gain of 2CLC features has already been very recently unveiled, highlighting a role for acetate, lactate and D-ribose ^32^. This metabolite screening also showed a positive effect of succinate on the 2C-like features acquisition. Beyond its role in the cellular metabolism, we show here that the succinate accumulation outside mitochondria leads to a global reduction in cytosine demethylation activity of the TETs and histone demethylation as previously observed in cancers after accumulation of succinate or SDH mutations ^54,55^. However, previous reports are somewhat discordant regarding the role of TETs in the acquisition of the 2CLC phenotype as their effect seems to be dependent on their interacting partners ^25,34,35,56^. For example, while TET2 could cooperates with PSPC1 (Paraspeckle Component 1) to reduce the expression of the retrotransposon MuERVL ^57^, binding of the TET proteins with SMCHD1 (structural maintenance of chromosomes flexible hinge domain containing 1) prevents the demethylation of the *Dux* gene locus and thus prevents its expression ^34^. Succinate accumulation would result in a global decrease in the activity of all 3 TET isoforms, resembling those of a TET triple KO, already shown to induce the 2C phenotype ^36^. The methylation landscape of histones, especially H3, is also highly dynamic both *in vivo*, at the time of the zygote genome activation (ZGA) that takes place at the 2C-stage, and in the 2CLC conversion with remodeling of H3K4me3, H3K9me3 and H3K27me3 ^58–60^ (extensively reviewed in ^61^). Similarly to the situation with the TETs, the action of HDM on the loss or acquisition of 2C-like features *in vitro* is complex, as loss of KDM1a (lysine demethylase 1a) is shown to promote the expression of *Zscan4* and *MUERVL* ^62^ whereas loss of KDM5a and b decreases the expression of the markers and blocks the ZGA *in vivo* ^63^. However, while the literature highlights a role for these modifications of the epigenetic landscape in the 2CLC emergence, this is not supported by the data presented here. This discrepancy could be due to differences in the culture conditions, as these previous studies use mESC grown in serum + LIF conditions, while we use ground naïve mESCs (in 2iL and serum-free culture). Such difference has already been previously described in the establishment of another pluripotent state of ESCs, the paused state^64^. Interestingly, instead of an epigenetic rewiring as the major cause of 2CLC reprogramming, our data shed light on succinate acting as a paracrine/autocrine signal, able to trigger the emergence of 2CLCs by an MCT-dependent export and the activation of SUCNR1 expressing cells. Further studies are now needed to precisely dissect which downstream actors is truly responsible for the activation of the 2C transcriptional program.

Our study thus brings an additional and different example of metabolic control of pluripotency, and adds succinate to previously reported metabolites such as S-adenosylmethionine (SAM) ^6^, alpha-ketoglutarate (αKG) ^65^, glutamine ^66^, acetyl-CoA ^67^, lactate or D-ribose^32^ that are known to regulate the pluripotent states mostly through their contribution to modifications of the epigenetic landscape (reviewed in ^68^). Further emphasis on the importance of succinate in early development is also demonstrated by the embryonic lethality of the SDH subunits in mice^69–72^. Altogether, these results indicate a critical role of both heme and succinate in the progression of the pluripotency continuum, ranging from the 2CLCs on one hand and to the acquisition of primed pluripotency on the other.

## Material and methods

### A. Cell culture

mESCs (ES-E14TG2a) were cultured in N2B27 medium consisting in 1:1 mixture of DMEM/F12 (Gibco, 31330-038) and Neurobasal Medium (Gibco, 21103-049) supplemented with 1x N-2 Supplement (Gibco, 17502-048), 1x B-27 Supplement (Gibco, 17504-044), 1/100 penicillin-streptomycin (Gibco, 15140-122), 1x MEM nonessential amino acids (NEAA) (Gibco, 11140-035), 1x GlutaMAX (Gibco, 35050-038), 1x sodium-pyruvate (Gibco, 11360-039) and 0.1 mM β-mercaptoethanol (Gibco, 31350-010). Naïve mESCs were maintained on 0.2 % gelatin (Sigma, G1393)-coated plates at a density of 50 000 cells/cm^2^ and in N2B27 medium complemented with 10^3^ U/ml of mLIF (ESGRO, ESG1107), 3 μM of GSK3 inhibitor (CHIR99021) (Peprotech, 2520691) and 1 μM of MEK inhibitor (PD0325901) (referred to as 2iL) (SelleckChem, S1036). Cells were passaged every 2-3 days using accutase (Stemcell Technologies, #07920). Cells were then collected by centrifugation at 1200 rpm for 3 minutes and counted before seeding. The transition to EpiSC was obtained by transferring naïve mESCs on 15 μg/ml fibronectin (Gibco, 33010-018)-coated plates at a density of 30 000 cells/cm^2^ and by supplementing the N2B27 medium with 12 ng/ml of bFGF (Peprotech, 100-18B) and 20 ng/ml of activin A (Peprotech, 120-14P). Coating proteins were incubated 1h before seeding. mESCs were maintained at 37°C, 5 % CO_2_ in a humidified incubator.

### B. mESC treatment

The heme inhibitor succinylacetone (SA) (Sigma, D1415) is used at a concentration of 0.5 mM. Hemin (Sigma, 51280) is used at a concentration of 10 μM in 0.1N NaOH. Diethyl butylmalonate (BM) (Sigma, 112038) is used at a concentration of 1 mM. Atpenin A5 (Santa Cruz biotechnology, sc-202475) was used at 250 nM. SIS3 (Selleck chemicals, S7959) was used at 5 μM. NF-56-EJ40 (Axon Medchem 3056) was used at 1μM. AZD3965 (Selleckchem S7339) was used at 1 μM. Exogenous succinate was provided as either 40 mM Sodium succinate (Sigma) or 16 mM dimethylsuccinate (Sigma) supplementation. HDM inhibition was achieved with 250 nM JIB04 (Medchem express HY-13953) and TET inhibition with 5 μM TETin-C35 (Aobious AOB11121).

### C. RNA extraction and RT-qPCR

RNA was extracted after 2 days of culture with the ReliaPrep™ RNA Tissue Miniprep System (Promega, Z6111) following manufacturer’s protocol for non-fibrous tissue by adding RNA lysis buffer on pelleted cells. RNA concentrations were quantified with the Nanophotometer N60 (Implen). Reverse transcription (RT) was performed with the GoScript™ Reverse Transcriptase kit Random Primers (Promega, A2801) to convert 1 μg of RNA into cDNA. Briefly, RNA was mixed with RNAse-free water to obtain 1 μg of RNA in 12 μL and heated 5 minutes at 70°C. Then, 8 μL of RT mix (4 μL random primers buffer, 2 μL enzyme, 2 μL RNAse free water) was added and the reaction was performed in a thermocycler (5 minutes at 25°C, 60 minutes at 20°C and 15 minutes at 70°C).

The qPCR was performed on the ViiA 7 Real-Time PCR System (ThermoFisher) with 10 ng of cDNA per reaction, SYBR Green GoTaq qPCR Master Mix (Promega, A6002) and primers listed in the **Table number 1** at a final concentration of 300 nM. Altogether, 2 μL of cDNA (5 ng/μL), 1 μL of forward primer (6 μM), 1 μL of reverse primer (6 μM), 10 μL of Master Mix and 6 μL of RNAse free water were added in each well. Relative expression was calculated using the 2^-ΔCt^ method with GAPDH as an endogenous control.

**Table 1:**
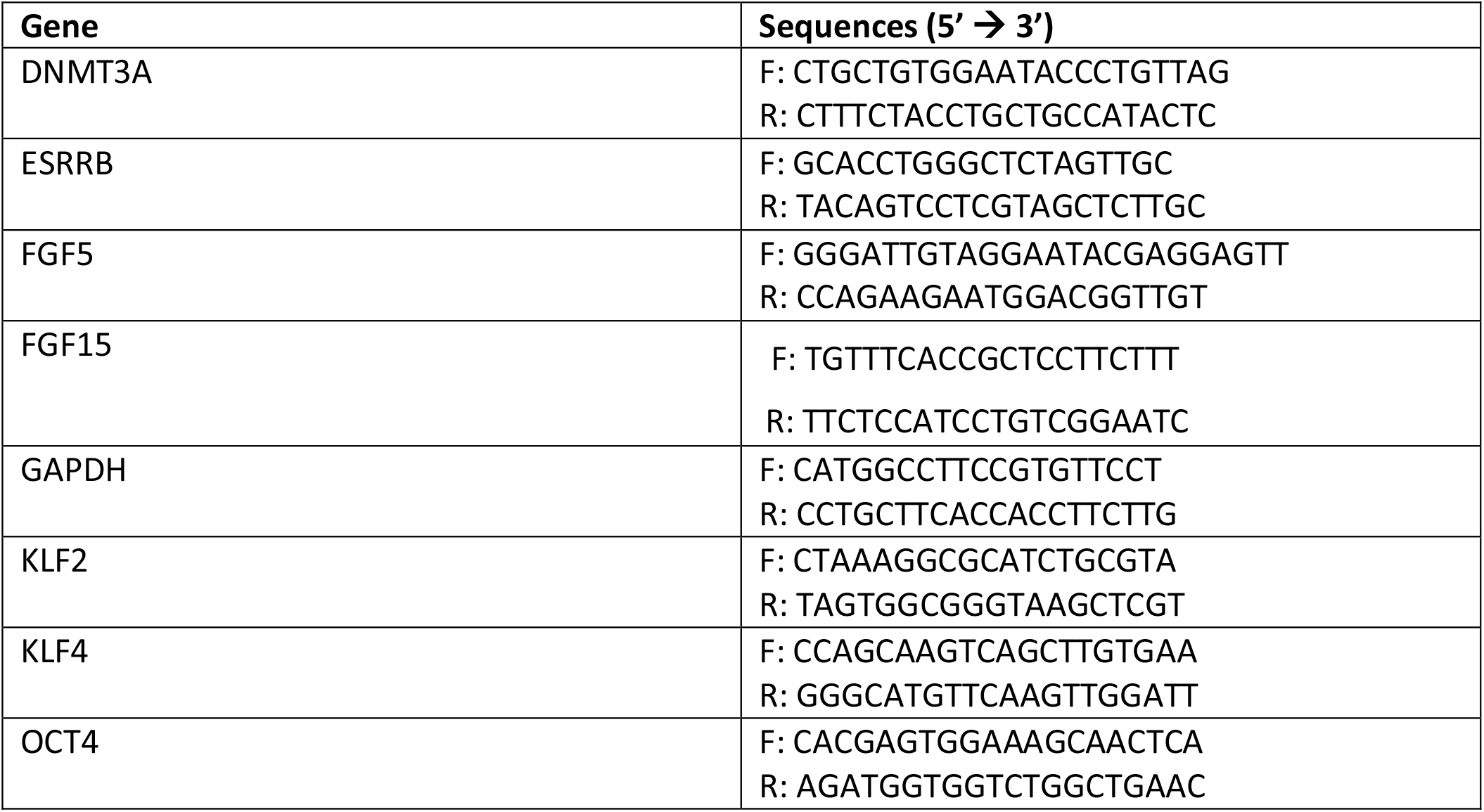
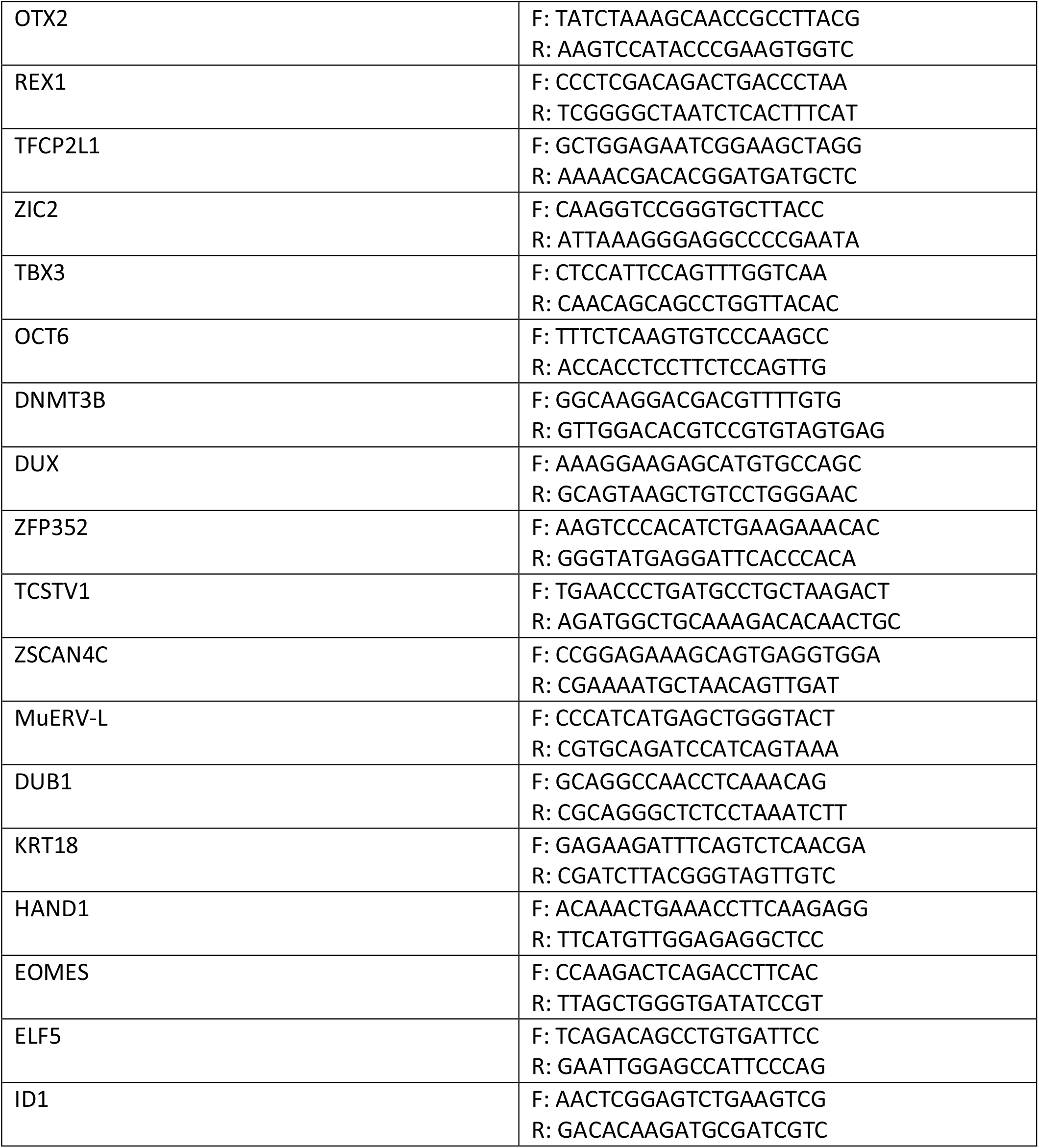
List of primers used in qPCR

### D. Western blot analyses

Pellets of cells were lysed by adding protein lysis buffer (20 mM Tris-HCl; pH 7.5, 150 mM NaCl, 15 % Glycerol, 2 % SDS, 25x protease inhibitor cocktail (PIC, cOmplete protease inhibitor cocktail, Roche 11697498001), 25x phosphatase inhibitor buffer (PIB, composed of 25 mM Na3VO3, 250 mM 4-nitrophenylphosphate, 250 mM β-glycerophosphate and 125 mM NaF), 1 % TRITON X-100, SuperNuclease (Sino Biologicals, 25 U/10μL) and by pipetting up and down. The protein concentration was determined by Pierce protein assay (ThermoFisher, 22660). Samples were mixed with Laemmli buffer (SDS, β-mercaptoethanol, Bromophenol blue) and heated for 5 min at 95°C before loading. An amount of 10 μg of proteins were loaded on SDS-containing 10 or 12 % polyacrylamide gels. At the end of migration, proteins were transferred to PVDF membranes (IPFL00010) by liquid transfer. Membranes were then blocked in LI-COR Intercept blocking buffer PBS for 1 h at RT and incubated overnight at 4 °C with the primary antibodies. Membranes were washed 3 times for 5 minutes with PBS + 0.1 % Tween-20 (PBST) before and after 1 h-incubation with the secondary antibodies at RT. Detections were performed and quantified with Odyssey LI-COR scanner. Primary and secondary antibodies were both diluted in Licor PBST. GAPDH was used as loading control. Primary and secondary antibodies for GAPDH detection were incubated 30 minutes. Antibody used are: anti-GAPDH (Sigma, G8795), anti-KLF4 (R&D, AF3158), anti-Oct4 (Santa Cruz, sc-5279), anti-OTX2 (R&D, AF1979), anti-pan-succinyllysines (PTM-401), anti-TFE3 (Sigma, HPA023881), anti-ZSCAN4 (Millipore, AB4340), anti-MUERVL-GAG (Novus, NBP2-66963), anti-ERK1/2 (Cell Signaling 9102), anti-phosphoERK1/2 (Cell Signaling 9101) and anti-SMAD2/3 (Cell Signaling 8685).

### E. Immunofluorescence

Cells were seeded on coated glass cover (Assistent) slips 2 days before fixation with 4 % paraformaldehyde (Sigma, 30525-89-4) for 15 minutes. Coating was performed for 1 h with 15 μg/ml of fibronectin or with 3.5 μg/cm^2^ Cell-Tak (VWR, 734-1081) diluted in 0.1 M sodium bicarbonate. Cells were permeabilized and blocked for 30 minutes incubation in blocking buffer (PBS, 0.1 % TRITON, 1 % BSA). Immunostaining was performed by an overnight incubation of cover slips at 4 °C on 30 μL drops containing primary antibody diluted in blocking buffer. After three washes of 5 minutes in blocking buffer, the cells were incubated for 1 h in the dark and at RT with 30 μL drops containing secondary antibody and DAPI (Sigma, 10 236 276 001) diluted 1:1000 in blocking buffer. Cover slips were mounted with Mowiol after three washes of 5 minutes in blocking buffer. Analyses were performed with a Leica TCS SP5 confocal microscope (Leica microsystems). For 5mC analysis, cells were fixed for 15 min with 4 % PFA, permeabilized for 10 min with PBS-0.5% Triton X-100 and treated with 4 N HCl for 20 min to denature DNA. Cells were then washed once with distilled water a 2h blocking with 2% BSA and 0.5% Triton X-100. After these steps, the staining process would resume as described above, using a 5mC primary antibody (Sigma SAB2702243) diluted 1:300 in blocking buffer (PBS, 0.1 % TRITON, 1 % BSA).

### F. Flow cytometry

Cells were detached by a 5 minutes incubation with Accutase and spun for 5 min at 400 g. The cell pellet was then washed with PBS-10 % FBS followed by another centrifugation. The pellet, containing at least 10^6^ cells, was fixed by adding 100 μl of 4 % PFA for 15 min. After centrifugation, the cells were permeabilized with the permeabilization buffer (Invitrogen) for 30 min at room temperature. Antibodies were incubated for 1h30 at room temperature in PBS-10 % FBS and briefly vortexed every 30 min. After three washes with PBS-10 % FBS, the cells were incubated with an anti-rabbit Alexa 647 1:1000 for 1 h. Cell pellets were washed twice before being resuspended in PBS-10 % FBS and analysed with the FACSVerse machine.

### G. RNA sequencing

Sequence libraries were prepared with the Lexogen QuantSeq 3’ mRNA-Seq library prep kit according to the manufacturer’s protocol. Samples were indexed to allow for multiplexing. Library quality and size range were assessed using a Bioanalyzer (Agilent Technologies) with the DNA 1000 kit (Agilent Technologies, California, USA). Libraries were subsequently sequenced on an Illumina HiSeq4000 instrument. Single-end reads of 50 bp length were produced with a minimum of 1 M reads per sample.

Quality control of raw reads was performed with FastQC v0.11.7, available online at: http://www.bioinformatics.babraham.ac.uk/projects/fastqc. Adapters were filtered with ea-utils fastq-mcf v1.05 (Erik Aronesty (2011), ea-utils: “Command-line tools for processing biological sequencing data”; https://github.com/ExpressionAnalysis/ea-utils). Splice-aware alignment was performed with HiSAT2 against the mouse reference genome mm10. Reads mapping to multiple loci in the reference genome were discarded. Resulting BAM files were handled with Samtools v1.5 ^73^. Quantification of reads per gene was performed with HT-seq Count v2.7.14. Count-based differential expression analysis was done with R-based Bioconductor package DESeq2. Reported p-values were adjusted for multiple testing with the Benjamini-Hochberg procedure, which controls false discovery rate (FDR).

### H. Data analysis

TMM normalized rLog transformed counts were used for Principal Component analysis using R package PCATools. Gene set enrichment analysis (GSEA) was made on gene list ranked on Log2FC using R package ClusterProfiler ^74^. For genes with FC>2 in MUERVL::Tomato^+^ list from ^14^, Z score was calculated from TMM-rLog transformed counts and plotted as heatmap using R package Heatmap.plus. Analysis was made using statistical programming language R.

### I. Statistical analysis

One-way ANOVA statistical test followed Turkey multiple comparison test or student T-Tests were conducted on all results when indicated using GraphPad Prism version 9.1.1, GraphPad Software, San Diego, California USA, www.graphpad.com.

### J. Data availability

RNA-Seq data: Gene Expression Omnibus GSE178089.

## Acknowledgement

We are grateful to Dr. Maria Helena Padilla-Torres (Institute of epigenetics and stem cells; Helmholtz Zentrum München) for kindly providing the 2C:::turboGFP mESC cell line. We also thank the Morphym platform – UNamur for the help with confocal and flow cytometry analysis.

This work was supported by the Fonds de la Recherche Scientifique – FNRS, 5, rue d’Egmont, 1000 Brussels. D.D. and M.C. are recipient of a (Fonds de la Recherche dans l’Industrie et l’Agriculture [FRIA], Belgium), fellowship and S.M. is recipient of a (Fonds National de la Recherche Scientifique [FNRS], Belgium) fellowship.

## Author contribution

Conceptualization: D.D and P.R.

Investigation: D.D., M.C., S.M., M.F. M.D. and L.F.

Writing – Original Draft: D.D. and P.R.

Writing – Review & Editing: D.D., M.C., S.M., P.R., T.A. and J.M

Funding Acquisition: D.D and P.R.

Supervision: P.R. and T.A.

---

**Suppl. Figure 1:**
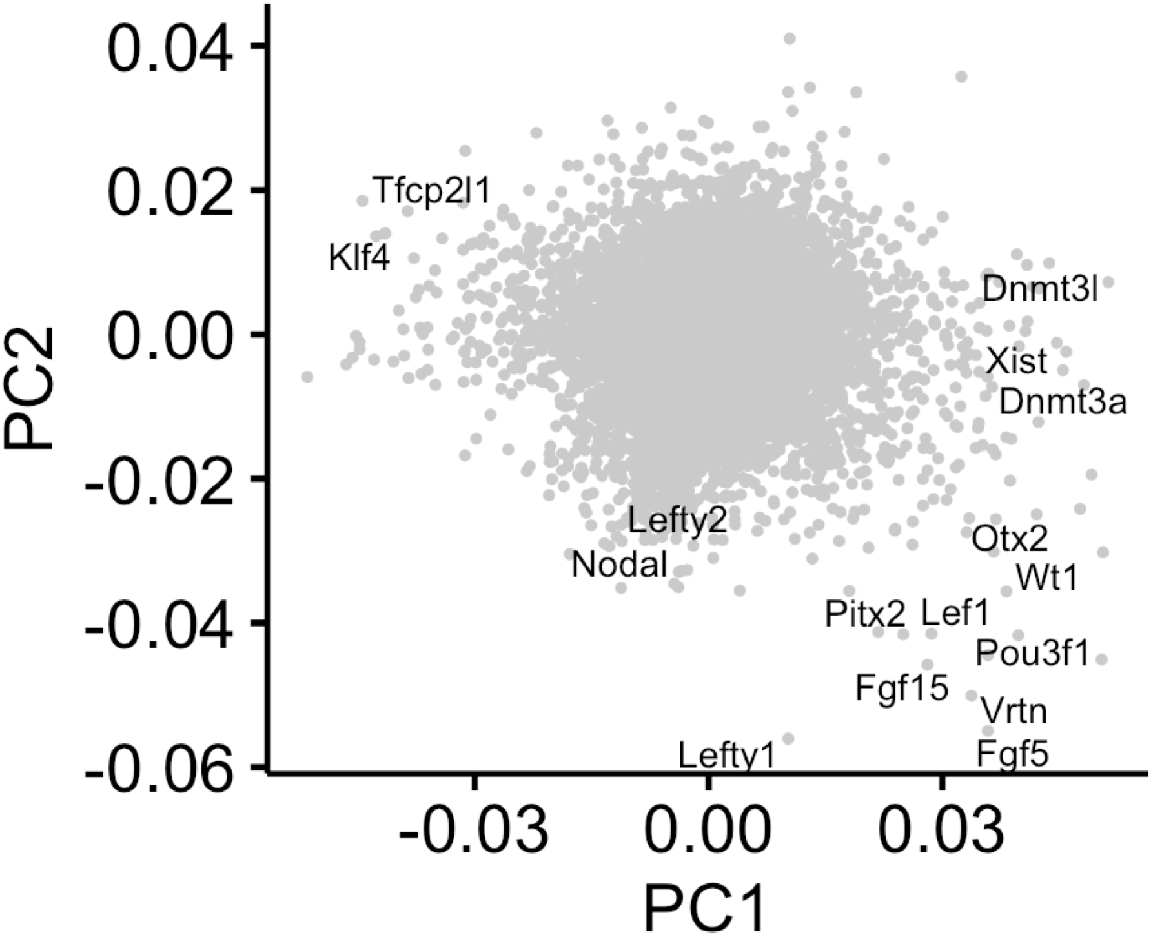
Relevant PC loadings. Genes contributing to principal components separating naïve 2iL, Epi with or without SA and hemin during the exit.

**Suppl. Figure 2:**
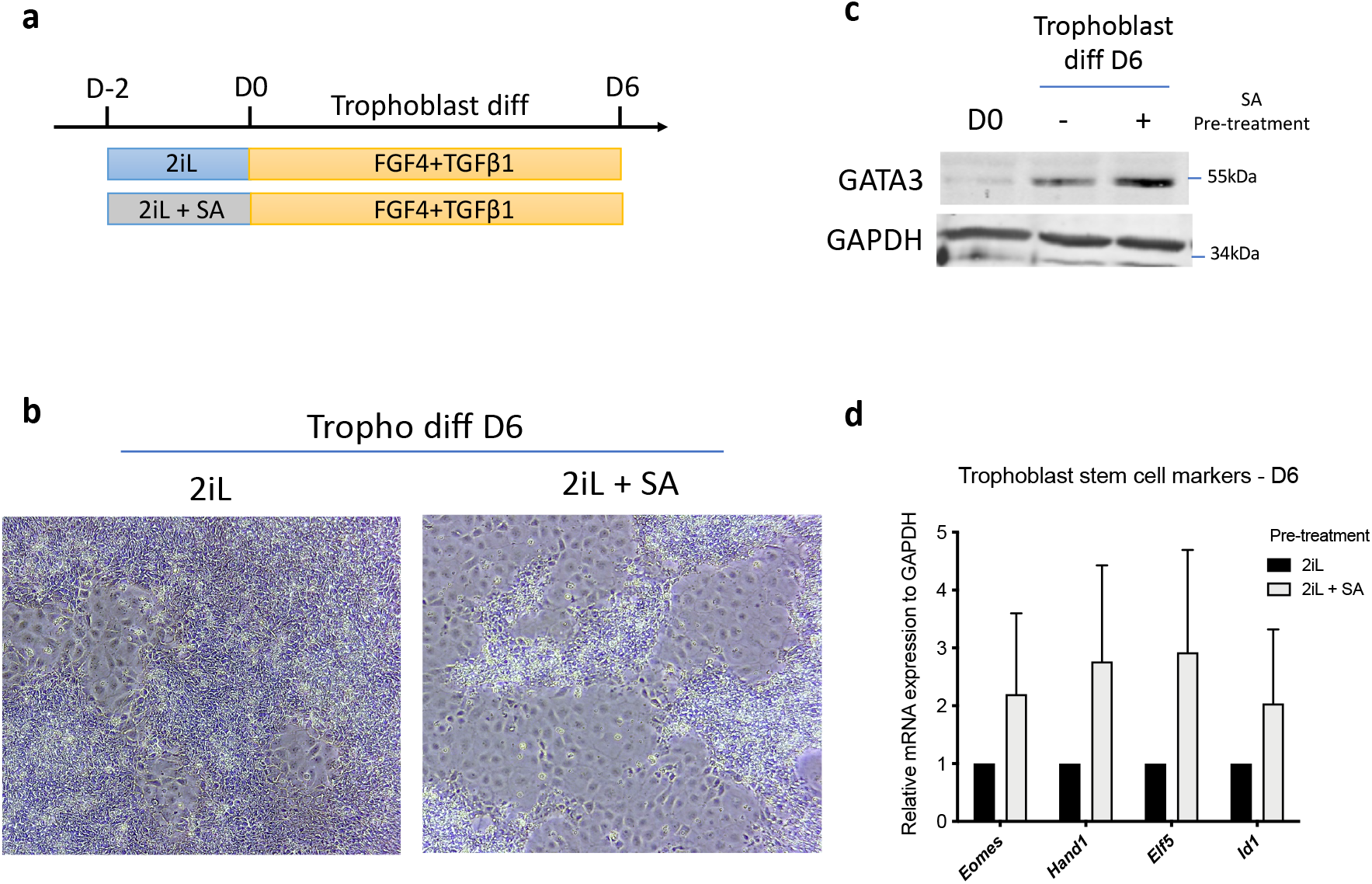
Pre-treatment with SA promotes a more efficient differentiation to trophoblast stem cells. **a)** Experimental diagram of the differentiation process of mESCs to trophoblast stem cells. Cells are first plated in naïve media with or without SA for two days before switching to a trophoblast differentiation medium (FGF4+TGFβ1) for 6 days. **b)** Phase contrast micrographs of cells after the differentiation process, pre-incubated or not with 0.5 mM SA. **c)** Western blot analysis of the protein abundance of GATA3 (GATA Binding Protein 3) relative to GAPDH loading control, in undifferentiated mESCs (D0) or after 6 days of trophoblast induction with or without heme synthesis inhibition (SA). Representative blot of 2 independent experiments. **d)** Relative expression of trophoblast stem cell markers of mESCs differentiated for 6 days with (2iL +SA) or without (2iL) pre-treatment with SA, assessed by RT-qPCR relative to *Gapdh* expression and normalized to 2iL condition. n=3 biological replicates. Pval > 0.05, not statistically significant

**Suppl. Figure 3:**
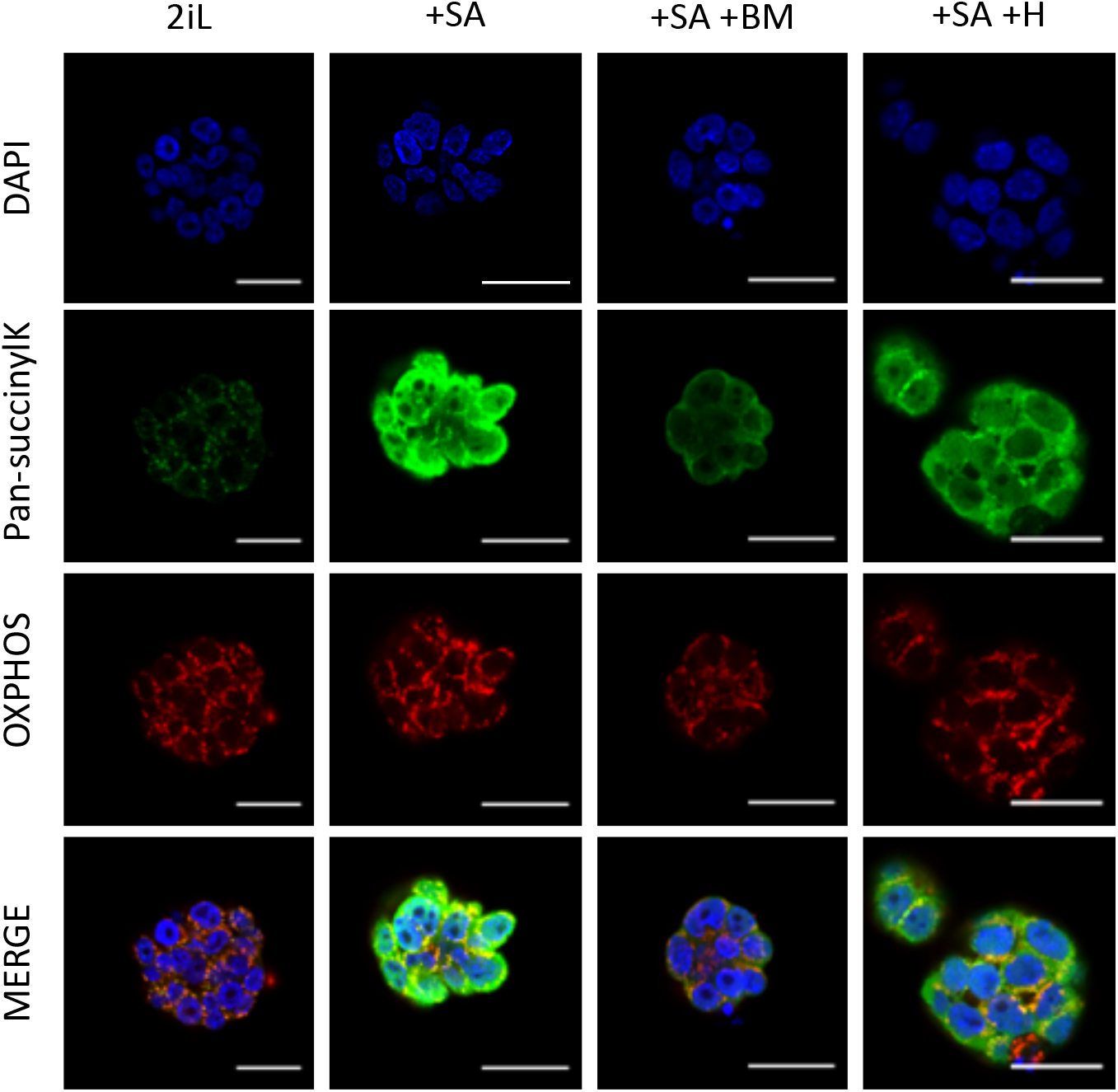
Treatment with SA induces a delocalization of succinyl-lysine-modified proteins outside of mitochondria. Immunostaining of succinyl-lysine residues (green) and OXPHOS complexes (red) in mESCs 2iL naïve control, in mESCs treated with 0.5 mM SA (+SA), with or without 10 μM Hemin (+SA + H), or 1 μM diethyl butylmalonate (+SA + BM). DAPI is used as nuclear counterstain. Representative images of n=3 independent experiments. Scale bar= 20 μm.

**Suppl. Figure 4:**
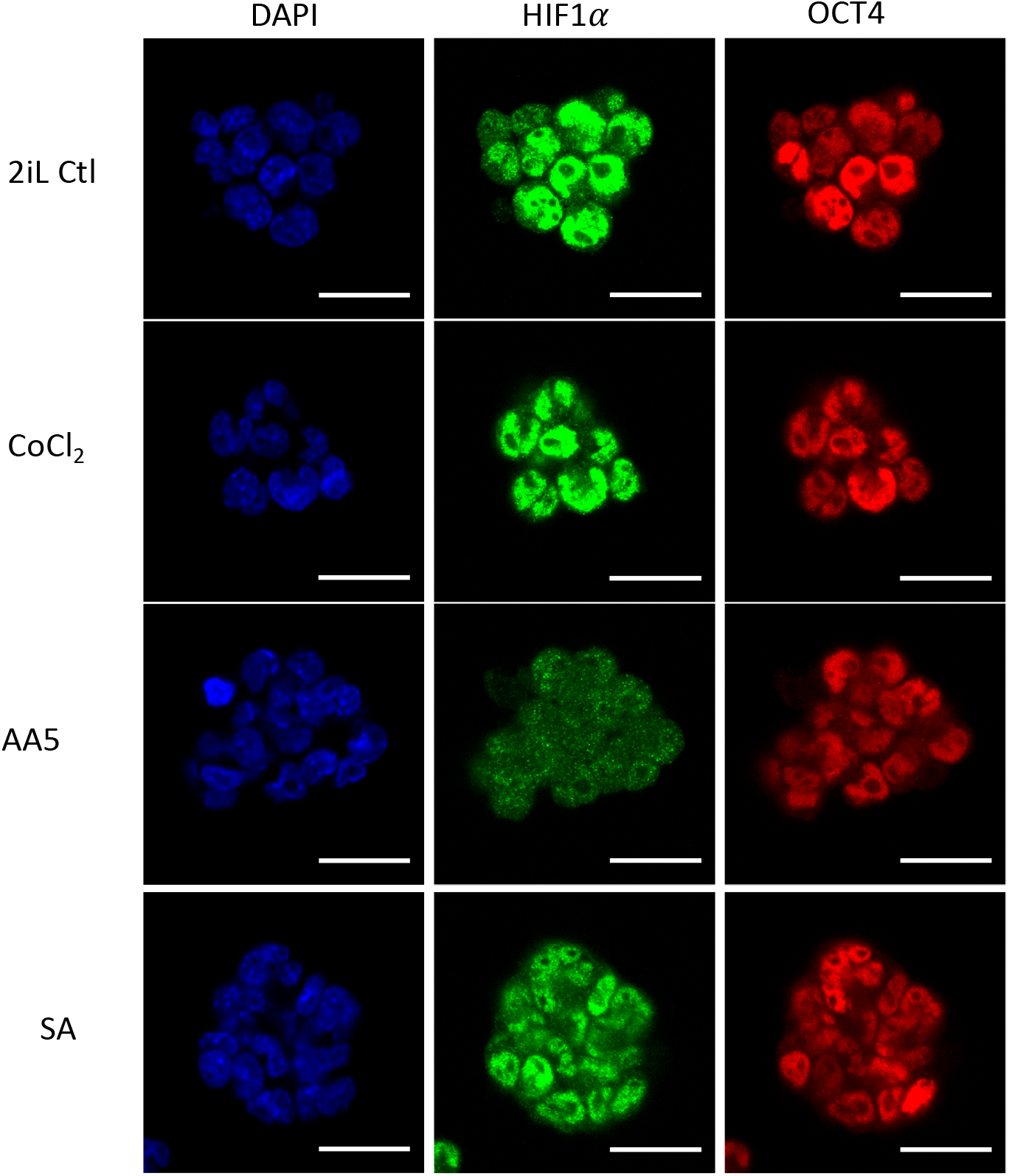
Treatment with SA or AA5 does not trigger HIF1*α* stabilization. Immunostaining of HIF1*α* (green), OCT4 as stemness marker (red) and DAPI as nuclear counterstain in mESCs 2iL naïve control, in mESCs treated with 100 μM of CoCl_2_ as positive control or 0.5 mM SA and 250 nM AA5. Representative images of n=3 independent experiments. Scale bar= 20μm.

**Suppl. Figure 5:**
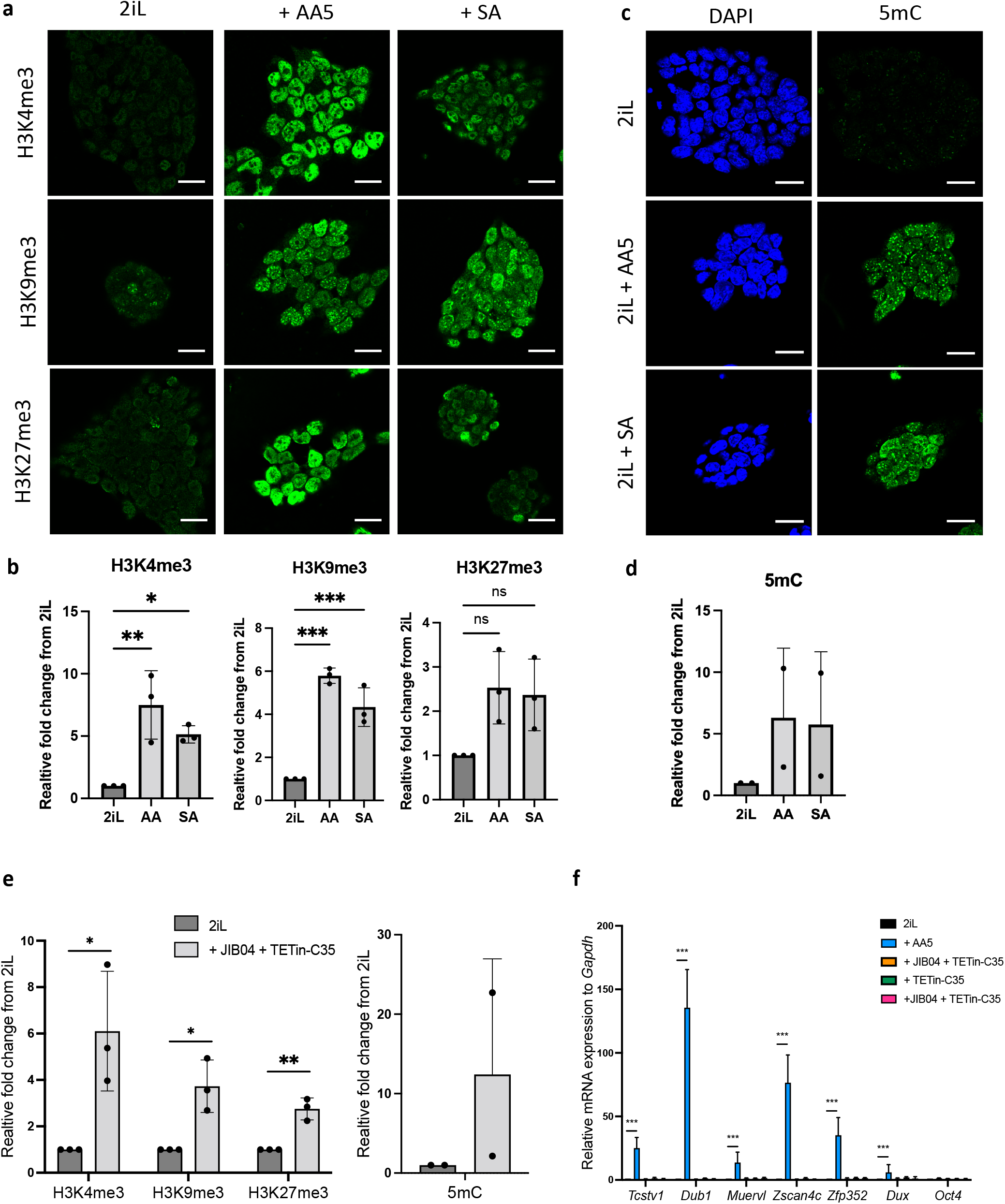
Treatment with SA or AA5 triggers an increase of the methylation of histones and DNA that does not cause a 2C-like reprogramming. a) Immunostaining of of trimethylated histone 3 (H3) residues (green) in mESCs 2iL naïve control, treated with 0.5 mM SA (+SA) or with 250nM AA5 (+AA5). Scale bar= 20 μm. b) Relative quantificaton of the methylated histone 3 fluorescent signal from confocal analysis. Data shown as mean +/-S.D. *p<0.05, **p < 0.01, ***p < 0.001. N.S. p>0.05 from ANOVA-1 analysis. n=3 independent biological replicates. c) Immunostaining of of 5-methylcytosine (5mC; green) in mESCs 2iL naïve control, treated with 0.5 mM SA (+SA) or with 250nM AA5 (+AA5). DAPI is used as nuclear counterstain. Scale bar= 20 μm. d) Relative quantification of the 5mC fluorescent signal from confocal analysis. Data shown as mean +/- S.D. n=2 independent biological replicates. e) Relative quantification of the trimethylated histone 3 (H3) residues signal from confocal analysis of naïve 2iL mESC or treated with 250 nM JIB04 as a HDM inhibitor and 5 µM TETin-C35 as a TET DNA demethylase inhibitor. Data shown as mean +/-S.D. n=3 independent biological replicates. T-tests f) Relative expression of 2C gene markers of mESCs assessed by RT-qPCR relative to *Gapdh* expression. Data shown as mean +/-S.D. *p<0.05, **p < 0.01, ***p < 0.001. T-test (e) or ANOVA-1 (f). n=3 biological replicates.

